# Three dimensional particle averaging for structural imaging of macromolecular complexes by localization microscopy

**DOI:** 10.1101/837575

**Authors:** Hamidreza Heydarian, Adrian Przybylski, Florian Schueder, Ralf Jungmann, Ben van Werkhoven, Jan Keller-Findeisen, Jonas Ries, Sjoerd Stallinga, Mark Bates, Bernd Rieger

## Abstract

We present an approach for 3D particle fusion in localization microscopy which dramatically increases signal-to-noise ratio and resolution in single particle analysis. Our method does not require a structural template, and properly handles anisotropic localization uncertainties. We demonstrate 3D particle reconstructions of the Nup107 subcomplex of the nuclear pore complex (NPC), cross-validated using multiple localization microscopy techniques, as well as two-color 3D reconstructions of the NPC, and reconstructions of DNA-origami tetrahedrons.

## Main text

Single molecule localization microscopy (SMLM) is capable of resolving biological structure at the nanometer scale. However, SMLM image resolution is ultimately limited by the density of the fluorescent labels on the structure of interest and the finite precision of each localization^1, 2^. Recently, methods for obtaining higher precision localizations have been reported, which work by either increasing the number of collected photons per molecule via e.g. cryogenic imaging^3, 4^, or by introducing patterned illumination^5, 6^. The first limitation remains, however, and one approach to boosting the apparent degree of labeling (DOL) and filling in missing labels can be applied when the sample consists of many identical copies of the structure of interest (e.g. a protein complex). In this case, by combining many structures into a single “super-particle”, the effective labelling density is increased, and the resulting super-particle has a high number of localizations leading to a significantly improved signal-to-noise ratio and resolution.

Previous approaches to this problem can be classified as either template-based or adaptations of existing single particle analysis (SPA) algorithms originally developed for cryo-electron microscopy (EM). Template-based methods^7, 8^ are computationally efficient, however, they are susceptible to template bias artefacts. Methods derived from SPA for cryo-EM have previously been adapted^9, 10^ and employed to generate 3D volumes from 2D projection data. These approaches are, however, intrinsically 2D to 3D, as they assume that the raw data are projections. Recently, Shi et. al^11^ also described a structure-specific method for 3D fusion, although they implicitly assume cylindrical particles and projected the volume onto top views only.

Here, we introduce a 3D particle fusion approach for SMLM which does not require, but can incorporate, a priori knowledge of the target structure. It works directly on 3D localizations, accounts for anisotropic localization uncertainties, and can perform cross-channel alignment of multi-color data. We demonstrate our method with 3D reconstructions of the Nuclear Pore Complex (NPC) obtained from three different SMLM techniques. The results exhibit a two orders of magnitude SNR amplification, and FSC-resolution values as low as 14-16 nm, which is sufficient to enable the identification of distinct proteins within a large macromolecular complex such as the NPC.

The processing pipeline is built upon our previous 2D method^12^ with modifications to each step to handle 3D localizations (**Figure 1a**). Briefly, we first register all *N* segmented particles in pairs, which provides *N*(*N* − 1)/2 relative registration parameters *M*_*ij*_ (3D rotation and translation from particle *i* to *j*). To find the absolute poses *M*_*i*_, we map the relative poses from the group of 3D rotations and translations, SE(3), to its associated Lie-algebra and then average them (**Online Methods**)^13^. With the absolute poses determined, we then recompute the relative transformations to perform a consistency check. This makes use of the geodesic distance on SO(3) between the initial relative rotations *R*_*ij*_ and the estimates from the Lie-algebraic averaging 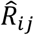:

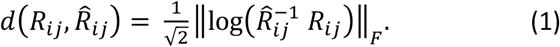

**Figure 1.**
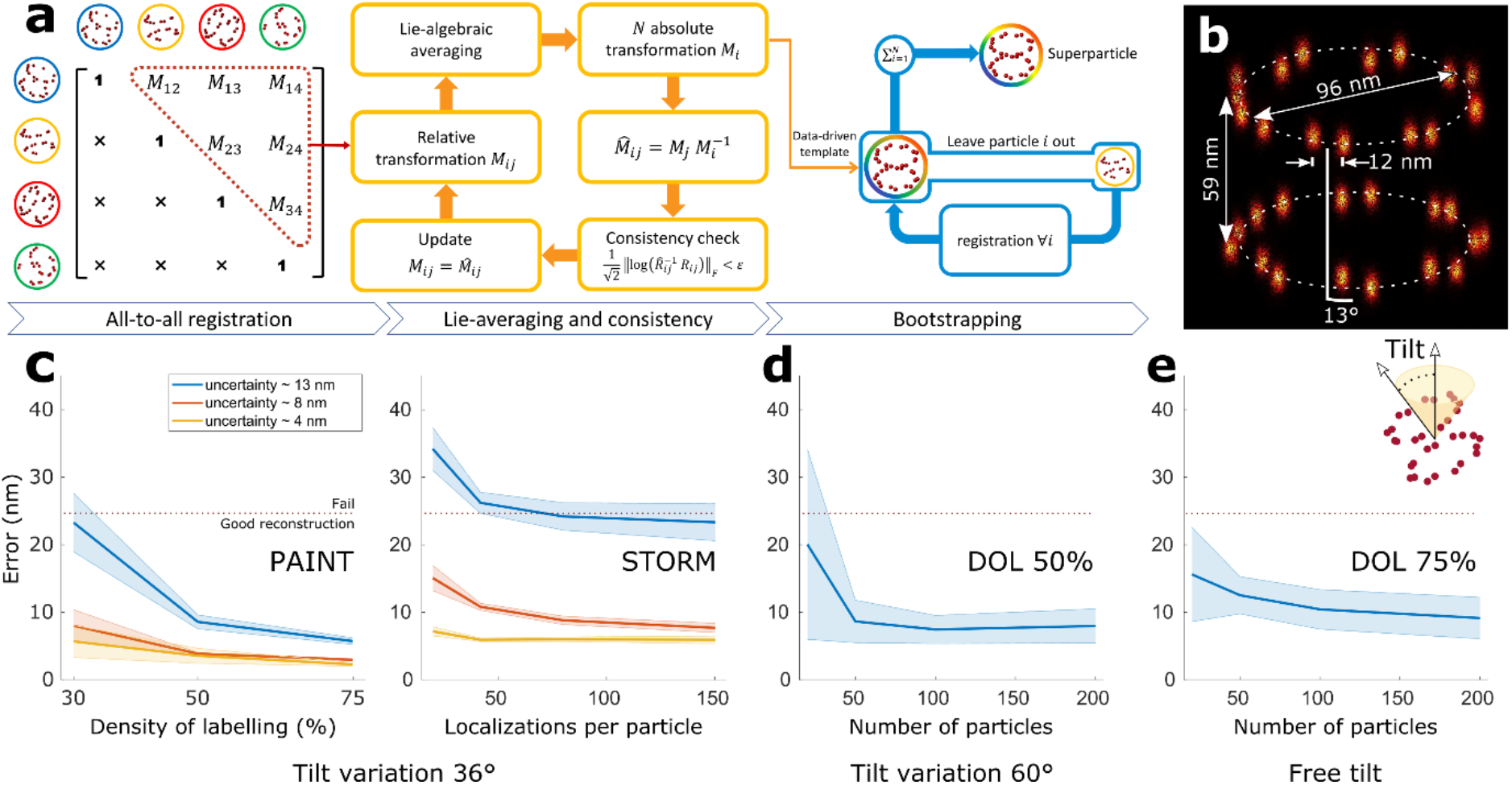
The 3D SMLM particle fusion pipeline and results of the simulation study. **(a)** Pair registration of all segmented particles results in relative transformations *M*_*ij*_ (translations and rotations). The redundant information in the *all-to-all registration* matrix is utilized for improving the registration errors by means of *Lie-algebraic averaging*, which results in *M_i_* absolute transformations. The relative transformations are recomputed as *M*_*j*_*M*_*i*_^−1^. From them, a *consistency check* is applied via a threshold ε on the rotation error to remove outlier registrations *M_ij_* from the all-to-all matrix. After two iterations, this results in a data-driven template. Finally, five rounds of *bootstrapping* are applied to improve the final reconstruction by registering every particle to the derived template. **(b)** Ground-truth fusion of 100 simulated NPCs indicating the height, radius, the angular shift between the cytoplasmic and nuclear rings in the same NPC. **(c)** Registration error for simulated PAINT and STORM data for different degree of labeling (DOL), mean localization uncertainties (σ = 4, 8 and 13 nm) and number of localizations per particle. Successful super-particle reconstruction is possible below a registration error of 25 nm. **(d)** Registration error of simulated PAINT data with 50% DOL and tilt angle of 60 degrees at different number of particles per dataset. **(e)** Registration error of simulated PAINT data with 75% DOL and arbitrary pose at different number of particles per dataset. Solid lines indicate the mean and shaded area show the standard error of the mean (n=15).

Here, log is the matrix logarithm and 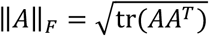 the Frobenius norm. The geodesic distance on SO(3) ranges between 0 and *π*, we set an empirical threshold ε = 1 radian to remove inconsistent pairwise registration entries in the all-to-all matrix. The transformations retained after the consistency check are used to generate a data-driven template. Each single particle is then registered to density-resampled versions of this template for 3-5 iterations. During this process, prior knowledge of symmetry can be incorporated (see **Online Methods**). Additionally, we propose a computationally efficient means of sorting and removing outliers (see **Online Methods**).

We evaluated our algorithm using simulations of the Nup107 subcomplex of the NPC (**Figure 1c-e**). Nup107 is a nucleoporin which is part of the Nup107-160 complex^14^ together with eight other nucleoporins. Our ground-truth model consists of 2×16 copies of Nup107 arranged in 8 pairs on the two rings of the NPC, with a 13° azimuthal shift (**Figure 1b**). The quality of registration was assessed with an error measure based on the residual registration error of the underlying binding sites (see **Online Methods**), which is independent of the localization precision. We found that for registration errors larger than the distance between the 8-fold symmetric subunits of the NPC rings (~25 nm) the reconstruction was so poor that we considered the alignment to be a failure (**Supplementary Figure 1**).

We simulated both PAINT and STORM imaging, to assess how the switching kinetics of the fluorescent labels affects the particle fusion (**Supplementary Figure 2**). For PAINT, we generated particles with a DOL of 75%, 50%, and 30%, localization uncertainties of 3, 8, and 13 nm in-plane and three times worse in the axial direction, and tilt angles spanning a range of +/−36 degrees (**Supplementary Figure 3**). For STORM, we kept the DOL fixed at a realistic value of 50% while varying the average number of localizations per particle from 20 to 150 (corresponding to different fluorophore bleaching rates), and with the same range of localization uncertainties and tilt angles as before. For each simulation condition, we generated 15 datasets containing 100 particles each. We found that a registration error below 8-10 nm was required (**Supplementary Figure 1**) to fully resolve the sixteen pairs of Nup107 sites. For PAINT, this was achieved for a minimum DOL of 50% and a localization precision better than 8 nm (**Figure 1c**). For STORM, we observe that for high localization precision (~4 nm) the registration error is below 10 nm even for a low number of localizations per particle (down to 20). For a lower average localization precision of ~13 nm, the registration errors of all simulated STORM datasets were above 20 nm. This is similar to the error range of PAINT data at 30% DOL. Consistent with our previous work therefore,^12^ we observe that STORM imaging requires a higher DOL than PAINT to achieve a similar performance. The simulations also indicate that a high-quality reconstruction (error <10 nm) requires at least 50-100 particles (**Figure 1d**) for PAINT data with 50% DOL. Even for unconstrained random pose variations and 75% DOL, the required number of particles for a successful registration remains relatively constant (**Figure 1e**).

We applied our algorithm to experimental SMLM data of NPCs in fixed U2OS cells (**Figure 2** & **Supplementary Movies 1-3**). Cells expressing Nup107-SNAP labeled with Alexa Fluor 647-benzylguanine were imaged with three different SMLM techniques, 3D astigmatic PAINT (**Supplementary Figure 5**), 3D astigmatic STORM^15, 16^ and 4Pi STORM^17, 18^. **Figure 2a, e** and **i** show the results of fusing 306, 356, and 750 segmented particles for the three modalities, which had an average number of localizations per particle of 88, 70, and 58, respectively. After fusion, the FSC resolution was ~15 nm (isotropic, see **Supplementary Figure 6**). We measured the distance between the cytoplasmic and nuclear rings as 60.5, 61.6 and 62.9 nm for PAINT, STORM and 4Pi STORM data, respectively (**Figure 2b, g** and **l**), and we determined the average radius to be 49.1, 53.2 and 51.1 nm and 50.8, 51.8, 52.8 nm for the two rings (**Figure 2c-d, h-i** and **m-n**). Finally, the phase shift differences between the two rings (for analysis see **Online Methods**) were found to be ~10°, 14° and 14° (**Figure 2e, j** and **o**, **Supplementary Figure 7**). These measurements are in good agreement with cryo-EM based models derived from the work of von Appen et al.^19^, who found a phase shift of 14°, height of 59 nm, outer ring radius of 49.7 nm and inner ring radius of 46.6 nm. The experiments for NPCs in the lower nuclear membrane indicate a narrow tilt angle distribution (~14°, see Supplementary Figure 4), well within the tilt tolerance limit assessed from the simulations.

**Figure 2.**
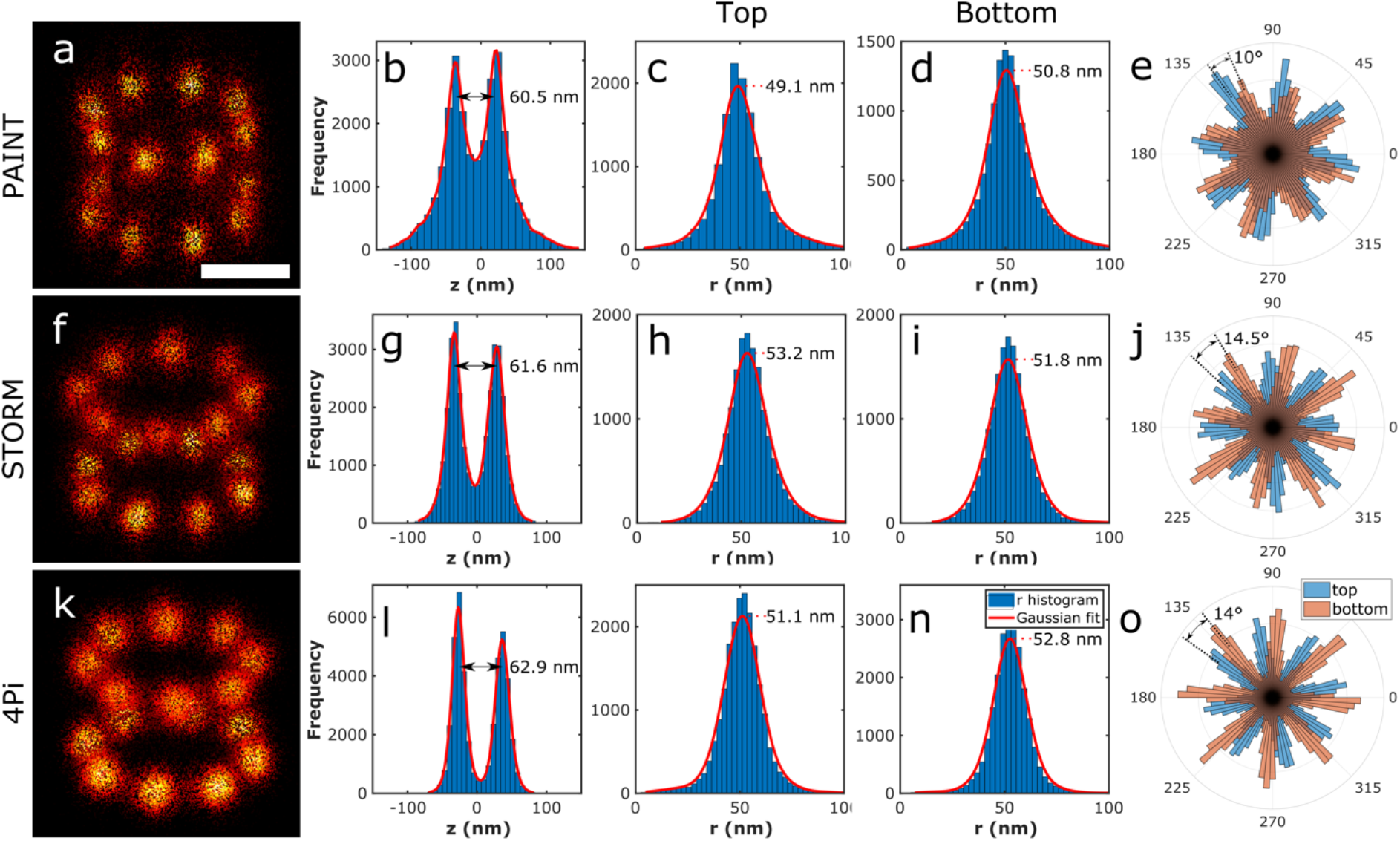
3D Particle fusion of Nup107 acquired with different 3D localization microscopy techniques. **(a)** Fusion of 306 particles acquire by 3D astigmatic PAINT. **(b)** Histogram of the Z coordinate of localizations in the super-particle. **(c)** Histogram of the radius of cytoplasmic ring localizations, **(d)** nuclear ring. **(e)** Rose plot of the localization distribution over azimuthal angles for the cytoplasmic (blue) and nuclear (orange) rings of the super-particle. **(f)** Fusion of 356 particles acquired by 3D astigmatic STORM. **(g-j)** Similar to **(b-e)**. **(k)** Fusion of 750 particles acquired by 4pi STORM. **(l-o)** Similar to **(b-e)**.

In a second experiment, we used multi-color 4Pi STORM to simultaneously visualize two components of the NPC (see **Figure 3**). U2OS cells expressing Nup107-SNAP were stained with Cy5.5-benzylguanine, and also with Wheat Germ Agglutinin (WGA) conjugated to Alexa Fluor 647, which is known to bind to FG-repeat nucleoporins in the central channel region of the NPC. First, we performed particle fusion on the Nup107 localizations. Next, we applied the transformations determined from the first step to the WGA channel, and then superimposed the two color channels in a single volume with a common origin. The resulting multi-color super-particle shows the location and dimensions of the central channel of the NPC with respect to the nuclear and cytoplasmic Nup107 rings. Despite the unstructured nature of the FG-repeats, by aligning with respect to the rigid Nup107 structure, the fused WGA data maps out the spatial distribution of FG-repeats within the channel.

**Figure 3.**
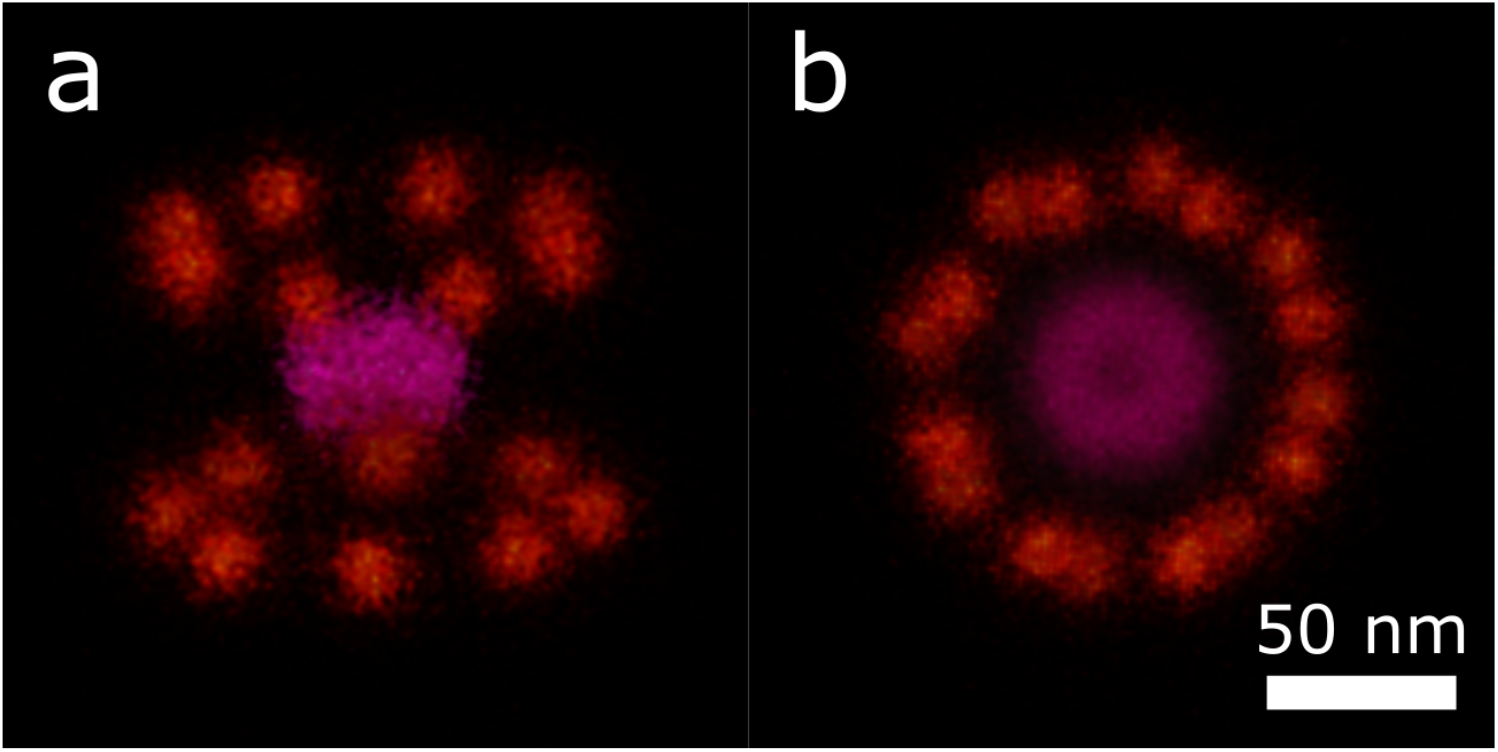
Fusion of 831 multi-colour 4pi STORM images of nuclear membranes stained for Nup107 and wheat germ agglutinin (WGA). **(a)** Side view. **(b)** Top view. Applying particle fusion on Nup107 channel (red) provides a set of absolute transformations which were subsequently used to align the corresponding WGA channel, which stains FG-repeat nucleoporins in the central channel region of the NPC (magenta).

In a final experiment, we fused 256 tetrahedron-shaped DNA origami nanostructures acquired with PAINT (**Supplementary Figure 8-10**). The height of the tetrahedron was measured from the peak-to-peak distance of the z-histogram of the super-particle to be ~90 nm (**Supplementary Figure 10d**). This implies a side length of 104 nm which agrees well with the origami design of 100 nm^20^.

In conclusion, we have developed a general purpose, template-free 3D particle fusion algorithm for SMLM that is robust to typical experimental conditions, and have shown its performance using simulations, the Nup107 subcomplex of the NPC for three different imaging setups, and DNA-origami tetrahedrons. By boosting the SNR of the data, our particle fusion method increases the effective spatial resolution and makes possible the reliable identification of protein locations within macromolecular complexes, thereby adding specificity to EM-SPA methods via correlative approaches. In addition, as few as 50 particles were required for accurate reconstructions, enabling the exciting possibility to detect transient, infrequently populated states.

## Author contributions

S.S., M.B. and B.R. conceived the project. H.H. wrote the original code in Matlab, performed simulations and analyzed data. A.P. and M.B. contributed to the Matlab code, performed simulations, and developed external bindings in C and binary files. B.v.W. wrote GPU code. J. K-F. wrote the external bindings for Python. F.S. and R.J. acquired 3D astigmatic PAINT data, J.R. acquired 3D astigmatic STORM data, and M.B. acquired (two-color) 4Pi STORM data. H.H., M.B., S.S. and B.R. wrote the paper, and all authors commented on the paper.

## Acknowledgements

We thank Avishek Chatterjee for giving Lie-algebra averaging code. We thank Yiming Li and Philipp Hoess for acquisition and analysis of the 3D astigmatic STORM data. The U2OS cell line was a kind gift of Jan Ellenberg.

## Methods

### Sample preparation

#### 1. 3D astigmatic PAINT

##### NUP107

###### Cell culture

U2-OS cells were passaged every other day and used between passage number 5 and 20. The cells were maintained in DMEM supplemented with 10 % Fetal Bovine Serum and 1 % Penicillin/Streptomycin. Passaging was performed using 1× PBS and Trypsin-EDTA 0.05 %. 24 h before immunostaining, cells were seeded on ibidi 8-well glass coverslips at 30,000 cells/well.

###### Cell fixation

Prefixation was performed with prewarmed 2.4 % Paraformaldehyde (PFA) for 20 seconds followed by the permeabilization at 0.4 % Trion-X 100 for 10 seconds. Next, cells were fixed (main fixation) with 2.4 % PFA for 30 min. After 3× rinsing with 1× PBS the cells were quenched with 50 mM Ammoniumchloride (in 1× PBS) for 4 minutes. Then, cells were washed 3× with 1×PBS followed by incubation in 1× PBS for 5 minutes twice. For SNAP-labeling, cells were incubated with 1 μM of SNAP-ligand-modified DNA oligomer in 0.5 % BSA and 1 mM DTT for 2 hours. Finally, cells were washed 3× for 5 min in 1× PBS, incubated with 1:1 dilution of 90 nm gold particles in 1× PBS as drift markers, washed 3 × 5 min and immediately imaged.

##### DNA origami tetrahedron

The tetrahedron DNA origami structures were formed in a one-pot reaction with a 50 μl total volume containing 10 nM scaffold strand (p8064), 100 nM core staples, 100 nM connector staples, 100 nM vertex staples, 100 nM biotin handles, 100 nM DNA-PAINT handles, and 1400 nM biotin anti-handles in folding buffer (1× TE (5 mM Tris, 1 mM EDTA) buffer with 10 mM MgCl_2_). The solution was annealed using a thermal ramp cooling from 80 to 4 °C over the course of 15 h. After self-assembly, the structures were mixed with 1× loading dye and then purified by agarose gel electrophoresis (1.5% agarose, 0.5× TAE, 10 mM MgCl_2_, 1× SYBR Safe) at 3 V/cm for 3 h. Gel bands were cut, crushed and filled into a Freeze’N Squeeze column and spun for 5 min at 1000×g at 4 °C.

#### 2. 3D astigmatic STORM

Samples and data for the STORM modality were prepared and acquired according to Li et al.^16^.

#### 3. 4PI STORM

##### Cell culture

The U2OS cells were seeded on 18 mm #1.5 round coverslips which had been sterilized in 70% ethanol, dried and washed three times with 1× PBS. All coverslips used for 4Pi-SMLM were coated with a mirror-reflective aluminum film over one quarter of their surface, for the purpose of alignment in the 4Pi microscope. Mirror coating was accomplished using a thermal evaporator at the Optics Workshop of the Max-Planck-Institute for Biophysical Chemistry, Göttingen. Seeded cells were allowed to attach overnight at 37°C and 5% CO2 in a cell culture incubator.

##### Cell fixation

Cells were rinsed twice with PBS and pre-fixed with 2,4% paraformaldehyde (PFA; Electron Microscopy Sciences; cat.# 15710) in PBS (+Ca2+/Mg2+) for 30 seconds. The cells were then immediately permeabilized with 0.5% Triton X-100 (Sigma-Aldrich; cat.# T8787) in PBS (+Ca2+/Mg2+) for 10 min and directly fixed afterwards with 2,4% paraformaldehyde (PFA; Electron Microscopy Sciences; cat.# 15710) in PBS (+Ca2+/Mg2+) for another 30 min.. After fixation, the samples were rinsed three times with PBS and quenched for remaining fixative with 50 mM NH4Cl for 5 min. After quenching, the sample was rinsed three times with PBS and washed three times for 5 min. with PBS. The fixed samples were immediately stained using one of the protocols described below.

##### NPC labeling with SNAP-tag

After fixation, samples were blocked with a few drops of Image-iT FX Signal Enhancer (Thermo-Fisher; cat.# I36933) for 30 min. The benzylguanine (BG)-conjugated AF647 (SNAP-Surface; NEB; cat.# S9136S) was diluted to 1 μM in blocking solution (0,5% (w/v) BSA, 1 mM DTT in 1× PBS) and incubated with the sample for 1 hour. This was followed by a final round of three rinsing and 5 min washing steps.

##### Dual-color NPC labeling

After fixation, samples were blocked with a few drops of Image-iT FX Signal Enhancer (Thermo-Fisher; cat.# I36933) for 30 min. Benzylguanine (BG)-conjugated Cy5.5 was synthesized and kindly provided by the Chemical Facility of the Max Plank Institute in Göttingen. The BG-Cy5.5 was diluted to 200 nM in blocking solution (0,5% (w/v) BSA, 1 mM DTT in 1× PBS) and incubated with the sample for 2 hours. Next, the sample was rinsed three times with PBS and washed three times for 5 min with PBS. Immediately prior to imaging, the samples were also stained with Wheat Germ Agglutinin (WGA) coupled to Alexa 647 (Thermo Fisher # W32466). First, the WGA-Alexa 647 was diluted in 1% BSA in PBS to a concentration of 0.04 ug/mL, and the sample was incubated in this solution for 5 minutes. The sample was then washed three times for 5 minutes with PBS.

### Single molecule experiments

#### 1. 3D astigmatic PAINT

##### NUP107

###### Setup

Fluorescence imaging was carried on an inverted microscope (Nikon Instruments, Eclipse Ti2) with the Perfect Focus System, applying an objective-type TIRF configuration with an oil-immersion objective (Nikon Instruments, Apo SR TIRF ×100, numerical aperture 1.49, Oil). A 561 nm (MPB Communications Inc., 2W, DPSS-system) laser was used for excitation. The laser beam was passed through cleanup filters (Chroma Technology, ZET561/10) and coupled into the microscope objective using a beam splitter (Chroma Technology, ZT561rdc). Fluorescence light was spectrally filtered with an emission filter (Chroma Technology, ET600/50 m and ET575lp) and imaged on a sCMOS camera (Andor, Zyla 4.2 Plus) without further magnification, resulting in an effective pixel size of 130 nm (after 2 × 2 binning).

###### Imaging

Imaging was carried out using an imager strand concentration of 1 nM (P3-Cy3B) in cell imaging buffer (buffer C) 30,000 frames were acquired at 200 ms integration time. The readout bandwidth was set to 200 MHz. Laser power (@561 nm) was set to 130 mW (measured before the back focal plane (BFP) of the objective), corresponding to 0.73 kW/cm^2^ at the sample plane.

###### Axial calibration

Calibration was carried out as described earlier^14^.

##### Tetrahedron

###### Setup

Tetrahedron imaging experiments were carried out on an inverted Nikon Eclipse Ti microscope (Nikon Instruments) with the Perfect Focus System, attached to a Yokogawa spinning disk unit (CSU-W1, Yokogawa Electric). An oil-immersion objective (Plan Apo 100×, NA 1.45, oil) was used for all experiments. The excitation laser (561 nm, 300 mW nominal, coherent sapphire or 532 nm, 400 mW nominal, Cobolt Samba) was directly coupled into the Yokogawa W1 unit using a lens (focal length f = 150 mm). The pinhole size of the disk was 50 μm. As dichroic mirror, a Di01-T405/488/568/647-13 × 15 × 0.5 from Semrock or t540spxxr-uf1 from Chroma was used. Fluorescence light was spectrally filtered with emission filters (607/36 nm from Semrock or ET585/65m + ET542lp from Chroma) and imaged on an EMCCD camera (iXon 897, Andor Technologies), resulting in a pixel size of 160 nm. The power at the objective was measured to be ~10% of the input power.

###### Imaging

For the tetrahedron imaging experiment (2 nM of P1-Cy3b imager in buffer B) the Andor iXon 897 with a readout bandwidth of 5 MHz at 16 bit and 5× pre-amp gain was used. The EM gain was set to 100. 30,000 frames with an integration time of 800 ms were acquired. Imaging was performed using the Yokogawa W1 spinning disk unit with an excitation intensity of ~226 W/cm^2^ at 561 nm at the sample (laser was set to ~38 mW). No additional magnification lens was used resulting in an effective pixel size of 160 nm.

###### Calibration

3D images were acquired using a plan-convex cylindrical lens with a focal length of f = 0.5 m, ~2 cm away from the camera chip. The calibration was done as in earlier studies. For the processing of the data the software package Picasso^21^ was used.

#### 2. Astigmatic 3D STORM

Samples and data for the STORM modality were prepared and acquired according to Li et al. ^16^. In short, homozygous Nup107-SNAP U2-OS cell lines were fixed and labeled with Alexa Fluor 647 – benzylguanine and imaged on a custom-built setup that contains a cylindrical lens in the emission path for astigmatic 3D localization. The data were fitted using an experimental PSF model calibrated using a z-stack of beads that were immobilized on the coverslip^15^. Subsequently, fitting errors induced by the refractive index mismatch were corrected based on a calibration of beads immobilized in a gel^16^.

#### 3. 4Pi STORM

##### Setup

The design of the 4Pi microscope was based on an earlier design published by Aquino et al.^17^, which was then extensively modified to achieve higher image quality and usability. Specifically, the design was changed to incorporate feedback systems to maintain the sample focus position, higher NA objectives to collect more light, a completely redesigned sample stage allowing for fast and reliable sample mounting and linear translation when adjusting the sample position, a redesigned 4Pi image cavity allowing for maintenance of the beam path alignment, and new acquisition and control software to allow accurate control of the instruments involved in the system stabilization and acquisition of the raw image data. The laser illumination sources used for STORM imaging included a red laser for imaging (642nm CW, 2W, MPB Communications Inc.) and a UV laser for molecule re-activation (405nm CW, 100mW, Coherent). Excitation light was controlled and modulated either directly via the laser controller or via an acousto-optic tunable filter (AA Opto Electronic). Variable angle TIRF or near-TIRF illumination was achieved by coupling all light sources through an optical fiber, whose output was positioned in an optical plane conjugate to the objective lens back focal plane. By placing the output of the fiber on a motorized translation stage, the illumination angle could be continuously varied for optimal signal to background ratio. The 4Pi microscope cavity was based on two high-NA objective lenses (Olympus, 100x, silicone oil immersion, NA 1.35). One objective was fixed in position on a mounting block while the other was adjustable in three dimensions using a 3-axis piezo stage (Physik Instrumente, P-733.3). The adjustable objective was also adjustable in tip/tilt and XYZ via micrometer screws for coarse positioning and alignment. Illumination and control beams were introduced into the 4Pi cavity and brought out again via dichroic mirrors (ZT405-488-561-640-950RPC, Chroma). The detected fluorescence from the two objectives was recombined at a 50:50 beam-splitter (Halle). Prior to the beam-splitter each detected beam passed through a quarter wave plate (Halle) and a custom Babinet-Soleil compensator made of quartz and BK7 glass, one of which with an adjustable thickness of quartz glass, which allowed a precise phase delay to be introduced between the P- and S-polarized fluorescence light. The remainder of the detection path consisted of an optical relay to crop and focus the overlaid P- and S-polarized images onto four quadrants of an EMCCD camera (Andor Ixon DU897) as previously described. Before the camera, the light was filtered with fluorescence emission filters (Semrock LP647RU, Semrock FF01-770SP) and optionally a dichroic mirror (Semrock FF685-Di02) which allowed the fluorescence in one polarization channel to be filtered selectively for two-color 4Pi-SMLM imaging. Control systems included the sample focus control and the objective alignment control, and each of these was based on an infra-red laser beam introduced into the 4Pi cavity. The sample focus control was based on a design similar to that used in a standard STORM microscope: an infrared beam (830nm laser diode, Thorlabs) was reflected from the sample-glass interface, and the position of the reflected beam was detected on a photodetector. Fine control of the sample position was maintained with a linear piezo stage (Physik Instrumente, P-752) mounted underneath the top section of the three-axis linear stage used for sample positioning (Newport, M-462-XYZ-M). For the objective alignment control, a second infra-red beam (940nm laser diode, Thorlabs) was collimated and passed through the two objective lenses, focusing at the sample plane. Any motion of the two objectives with respect to each other resulted in a lateral shift in the transmitted beam, or a change in the collimation of the transmitted beam. The lateral shift was continuously monitored via a quadrant photodiode, and the transmitted beam collimation was monitored by splitting the beam and focusing it onto two pinholes positioned on either side of the focus, with photodetectors behind each pinhole. These signals were measured using a DAQ card (National Instruments), and a software-based feedback loop was then used to adjust the position of the movable objective lens to keep it aligned with the fixed objective lens. All microscope control and data acquisition were performed using custom software written in Labview (National Instruments).

##### Imaging

The sample was illuminated with 642 nm excitation light in order to switch off the fluorophores and cause them to blink stochastically. The emitted light was filtered spectrally (see above) and detected at the EMCCD camera, running at a frame rate of 101 Hz. Typically, 100000 image frames were acquired in a single measurement. During the experiment, the power of the 405 nm laser was manually adjusted to re-activate the fluorophores and keep the number of localizations per frame constant. Optical stabilization of the z-focus (focus-lock) was engaged before starting each recording, in order to minimize sample drift during the measurement. Prior to each set of 4Pi measurements, images of a fluorescent bead located on the sample were recorded as the bead was scanned in the Z-dimension, in order to create a calibration scan which was used in post-processing analysis of the 4Pi STORM image data. For all experiments, images of beads located at different positions in the sample plane were recorded, in order to generate a coordinate mapping which allowed the coordinate systems of the different image channels to be mapped onto each other.

##### Image reconstruction

STORM image analysis and reconstruction follows a standard approach based on peak finding and localization^22^. Two color imaging via the ratiometric method was analyzed as described previously^17, 23^. Correction of sample drift in post-processing was done based on image correlation of the 3D STORM data with itself over multiple time windows. STORM images were rendered as summed Gaussian peaks with a Gaussian width approximately equal to the previously measured localization precision (typically 3.5 nm in X, Y, and Z).

### Data fusion pipeline

Our data fusion framework is largely the same as our earlier work^12^ with 3D instead of 2D localization data. The anisotropic localization precision in 3D is naturally incorporated into the pair-wise alignment procedure using the Bhattacharya distance. We have to replace the consistency evaluation as rotations in 2D can be characterized by one in-plane angle only and therefore a straightforward threshold can be applied to the angle difference. In 3D, the three Eulerian angles are required to describe a rotation which complicates matter significantly as different rotations do not commute. To this end we used the geodesic distance eq. (1) on SO(3) as a measure for the dissimilarity between different rotations. Next to this necessary change for applying the framework in 3D, we have also made two other modifications to the earlier pipeline.

#### 1. Incorporation of symmetry

For symmetric structures and in the case of underlabeling or a non-uniform distribution of localizations per binding sites (e.g. in STORM), the hotspot problem reported earlier^12^ is unavoidable. The registration always tries to match dense regions of the structure and consequently the unbalanced occupancy of sites is reinforced in the process. We overcome this problem by properly incorporating prior knowledge about the symmetry group of the structure. For NPC, which has an eight-fold rotational symmetry (2D cyclic group *C*_8_) around the rotation axis through the center of the cytoplasmic and nuclear rings, we randomly added integer multiples of 2*π*/8 to the alignment angles of the particles at each iteration of the bootstrapping. This subsequently results in a uniform distribution of localizations over the binding sites. It is worth mentioning that this approach is different than what is done in single particle averaging in EM^24^ and in the method of Sieben et al.^10^ where the asymmetrical subunit of the particles is replicated to generate a symmetric structure based on the given symmetry group. In our approach the final reconstruction is mathematically *not* symmetric, but the symmetry is used to resolve the hotspot problem. This approach can easily be adapted to other simple point groups such as cyclic *C*_*n*_ and dihedral *D*_*n*_ groups given the axis (or axes) of rotation(s).

#### 2. Outlier particle removal

In our earlier work^12^, we kept all initially picked particles for the final super-particle. We only removed many of the bad registrations from the all-to-all matrix as long as the graph stays connected. In practice, however, it happens that the segmented particle set contains “outliers” that are either not a particle but background or just very low-quality particles. We propose a simple and efficient method for excluding outliers with small computational cost. After the bootstrapping step, we construct an *N* × *N* matrix with elements equal to the Bhattacharya cost function for all pairs of aligned particles (**Supplementary Figure 11a**). We sum over the columns (or rows) of this similarity matrix to assign a single score to each individual particle. If all particles are of good quality, these scores should be similar in magnitude. For outlier particles, however, we observe that the histogram of scores has an extended tail. We identify outliers as particles with scores are more than three scaled median absolute deviations (MAD) away from the median (**Supplementary Figure 11b**). This outlier particle removal only works properly if most of the segmented particles are of good quality and the particle fusion has not failed. The visual experience of the final reconstruction is barely affected for the examples shown in **Figure 2**, however, the best and worst particles demonstrate how this approach can rank the quality of the included particles (**Supplementary Figure 11c-d**).

##### Simulation setup

Our ground-truth model consists of 2×16 copies of NUP107 arranged in 8 pairs on the cytoplasmic and nuclear ring of the NPC with ~13 degree of azimuthal shift (**Figure 1b**). PAINT and STORM switching kinetics were simulated as earlier described^12^. For each parameter setting, we generated 15 datasets containing 100 particles each.

##### Registration error measure in simulations

To assess the performance of the method on simulated data, we devised an error metric which is independent of the shape of the ground-truth super-particle, does not have a global offset problem i.e. any transformation of the whole ensemble of particles gives the same error, can solve the symmetry ambiguity, is not impaired by underlabelling and has the same unit as the localization data. The error is the averaged Euclidean distance between corresponding binding sites after applying the data fusion process. This works in simulation only as there we know the ground truth and thus, we can establish point-corresponding between binding sites. This measures the registration error, however, if we would do the same with the localization data, we would get a convoluted compound of registration error and localization error and an overweighting of sites with many localizations. In **Supplementary Figure 12** **&** **13**, we illustrate the process. We find the point correspondence by measuring the distance for all possible combinations of binding sites and then report the minimum as the registration error between the two particles. **Supplementary Figure 13** demonstrates such combinations for a simplified NUP structure with *K* = 16 designed binding sites. Mathematically, the registration error of *N* aligned particles is computed as follows:

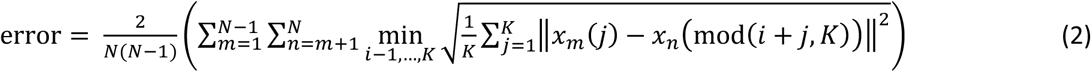

in which *x*_*m*_ is the localization data (3D coordinate) of the particle *m* from the set of all particles and mod is the modulo operator. The double sum sums over all pairs of particles and the sum over all possible correspondence of the binding sites for the current pair of particles.

##### Pose variation in simulation

In simulation we allow full orientationally freedom, which is not encountered our experiment. Due to the linear approximation of the exponential mapping from SO(3) to the Lie-algebra representation, averaging on SE(3) works only if the pose variation of particles is within a certain range. Therefore, fusion of particles with arbitrary poses can result in clusters of particles which are aligned within clusters but not across them (**Supplementary Figure 14a-b**). We developed a work-around for this problem as follows. As for outlier removal routine, we first compute the similarity matrix. Then, we subtract each row (or column) from the self-similarity (the Bhattacharya distance of a particle from itself) of the corresponding particle to convert the matrix into a dissimilarity matrix. We, then use multidimensional scaling (MDS)^25^ to translate the information about the pairwise distances between the *N* particles into a constellation of *N* points in Cartesian two-dimensional space. Subsequently, we use k-means clustering to identify clusters of particles (**Supplementary Figure 14c**). The user can easily identify the number of clusters in the MDS plot and for the current experiments we set it empirically to 3-4. Since the particles within each cluster are already well aligned (**Supplementary Figure 14d-f**), one can do a pairwise registration at the end to align all or some of the identified clusters.

##### Analysis of NPC structural parameters

NPCs are embedded in the nuclear membrane and their tilt axis aligns reasonably with the optical axis (normal distribution with about zero mean, **Supplementary Figure 4**). Consequently, the Lie-algebra always aligns the particles with the x-y plane for experimental data. A moment analysis of the super-particle is used to align the average pose with the principle planes (xy,xz, yz and etc.), i.e. aligning the symmetry axis of the NPC super-particle with the z-axis. The distance between the upper and lower rings of the NPCs is estimated by first computing the histogram of the z coordinate of the localization data in the super-particle. Then, a kernel-smoothing distribution with a bandwidth of 4 nm is fitted to the histogram and, finally, the distance between the two peaks of the fit is computed (**Figure 2b, g** and **l**). The radius of the two rings is measured by separating the localization data of the super-particle in two halves using a segmentation threshold which is computed as the local minimum of the z coordinate histogram. Then, the x and y coordinates of the localization data are transformed to two-dimensional polar coordinates (r, θ). The peak of the histogram of the r component of the localizations defines the radius of the rings (**Figure 2c-d, h-i** and **m-n**). The angular shift between the two rings of the NUP107 is estimated by, first fitting the function *b*_0_ + *b*_2_sin (8*θ* + *b*_2_) to the angular components of the localization data in each ring. The iterative least squares method is used for this nonlinear regression model to find the unknown coefficients *b*_0_, *b*_1_ and *b*_2_. Then, the difference between the fitted *b*_2_ parameters for the two rings defines the angular phase difference (**Supplementary Figure 7**).

---

**Supplementary Figure 1.**
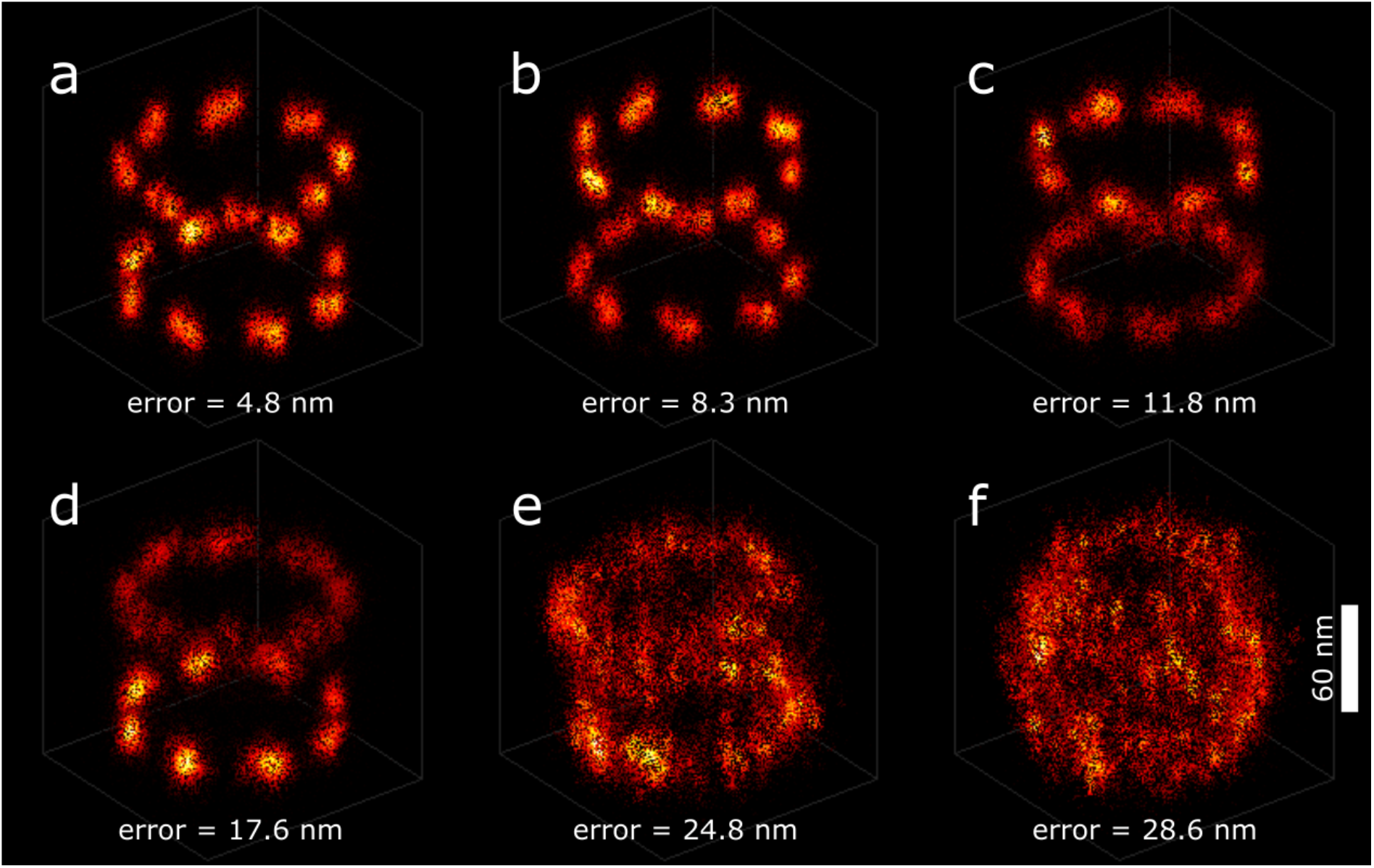
Particle fusion error for different reconstructions of a simulated STORM dataset. **(a-f)** Each reconstruction is the result of fusing 100 simulated particles with an average localization uncertainty of 4 nm and 50% DOL. From top to bottom and left to right, the error is increasing which also visually matches the quality of the reconstructions. For the error less than ~10 nm (**a-b**), the double blobs are still recognizable. For larger error as in (**c-d**), the double blobs merge into a single blob and for errors above ~20 nm the reconstructions lose the geometrical features of the ground-truth.

**Supplementary Figure 2.**
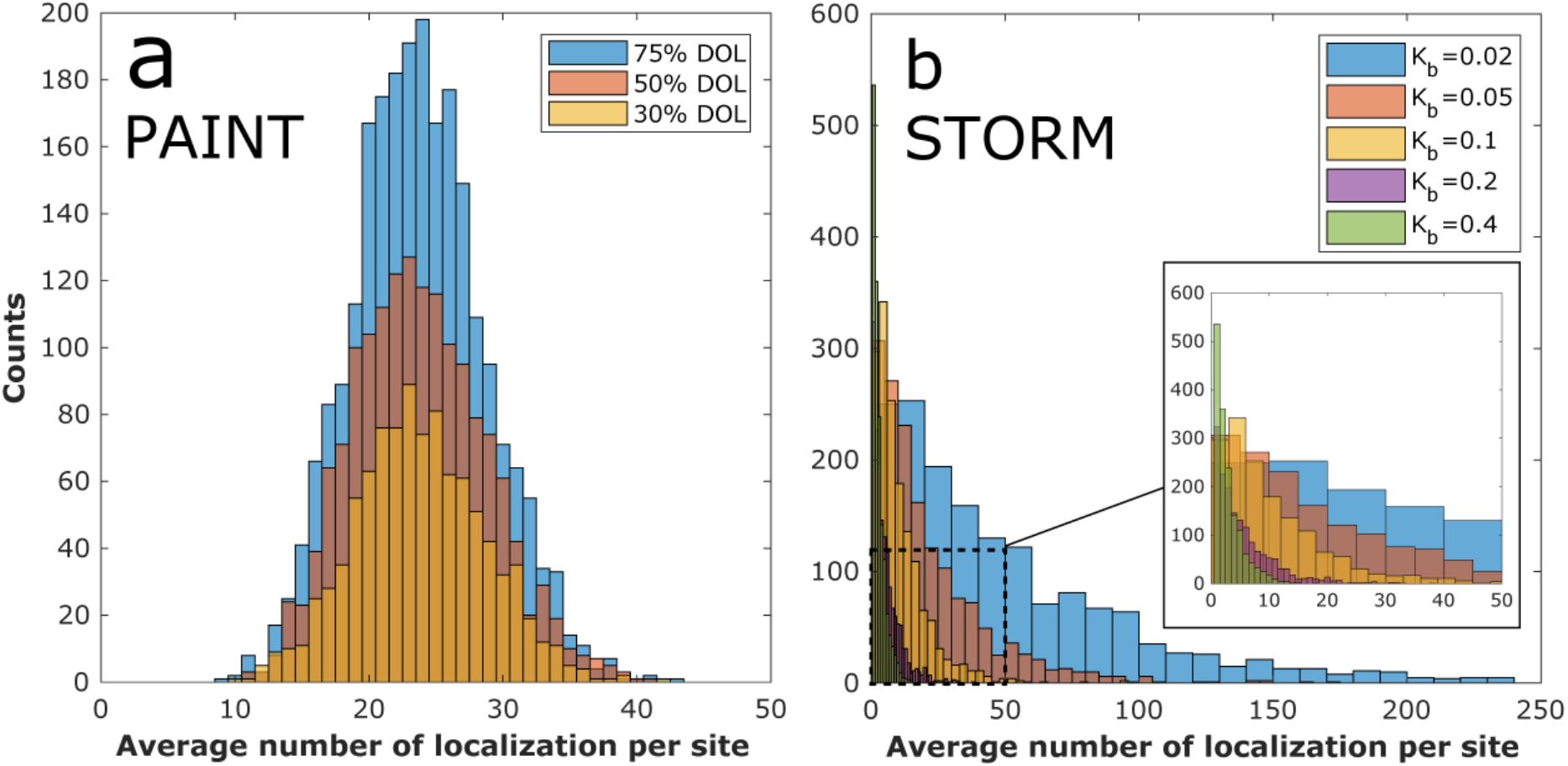
Statistics of localizations per binding sites for **(a)** PAINT and (**b**) STORM simulations. For PAINT particles, the distribution of localizations per site follows a Gaussian distribution while for STORM it is a Poisson. In case of STORM data, higher bleaching rate result in fewer localization per sites and a narrower bandwidth for the distribution.

**Supplementary Figure 3.**
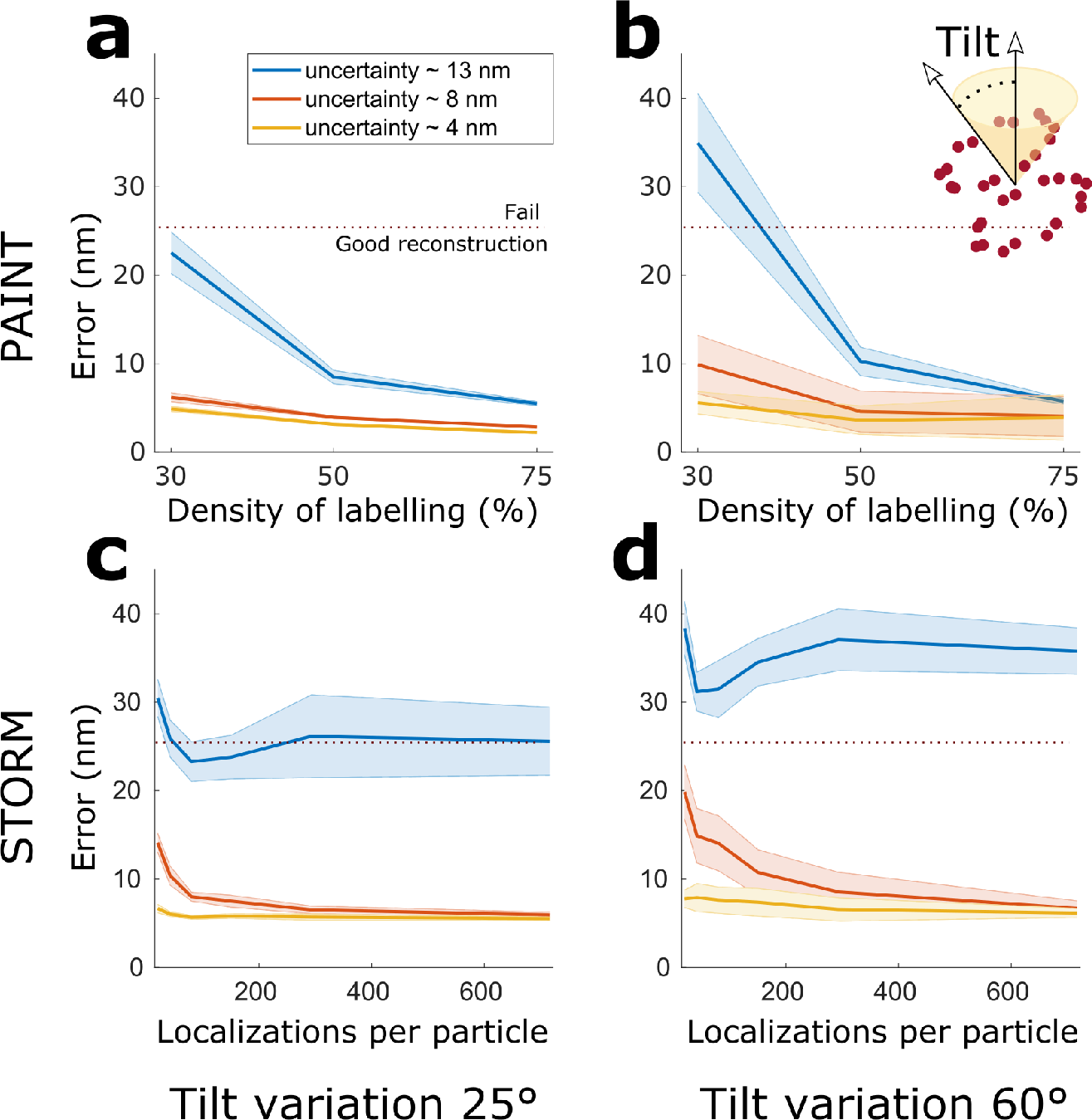
Particle fusion error for the alignment of PAINT and STORM SMLM images of simulated NUP107 particles with different initial tilt variations. **(a-b)** Particle fusion error of simulated PAINT data for different DOLs and for two range of tilt variations. **(c-d)** Particle fusion error of simulated STORM data for different number of localizations per particle (proportional to bleaching rate) and for two range of tilt variations. The particle fusion performance is getting slightly worse by increasing the tilt variations but in general it is quite stable even at high tilt angle range of 60 degree

**Supplementary Figure 4.**
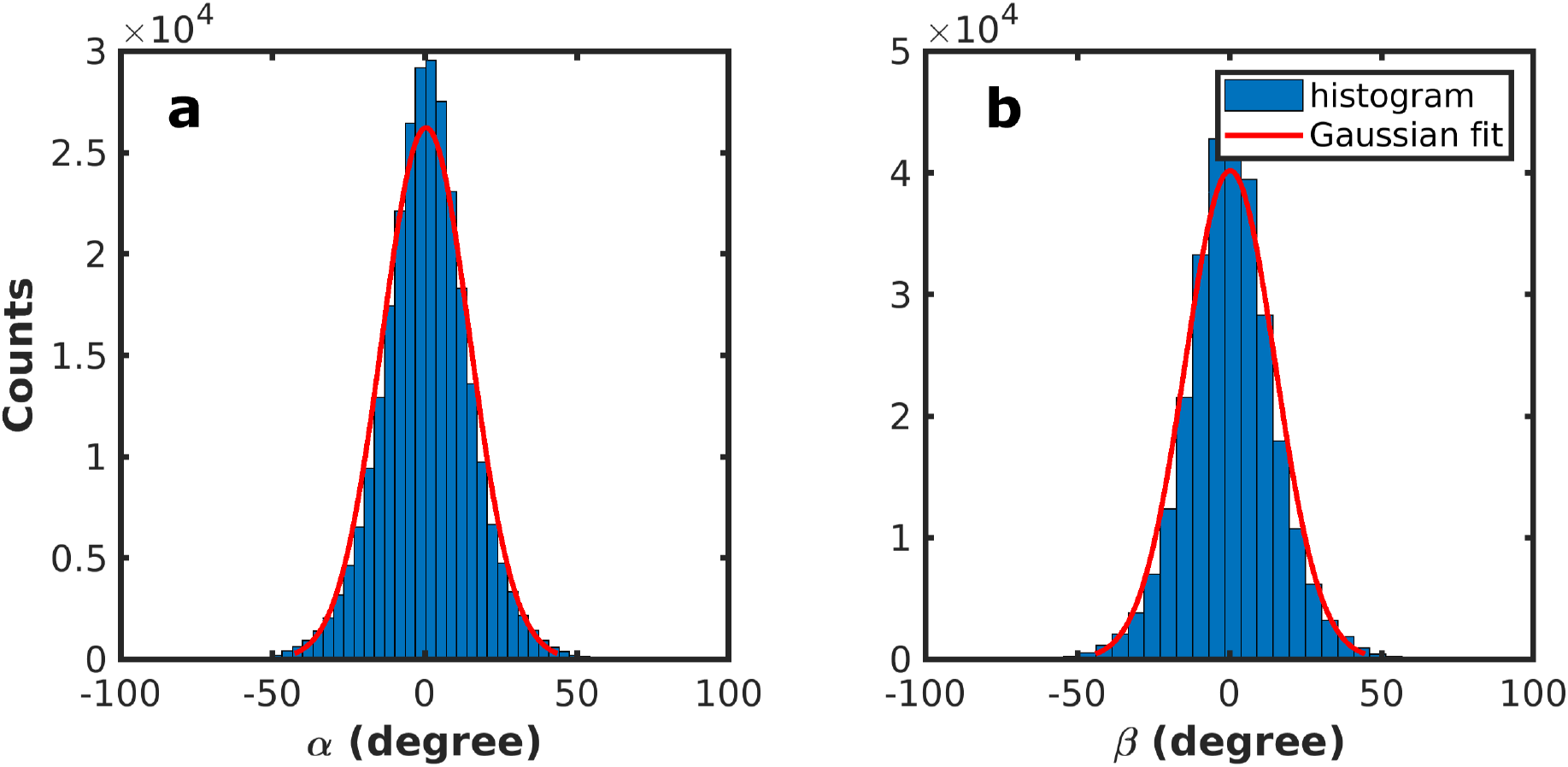
Tilt variations of the 4Pi experimental particles. **(a-b)** The histograms of the Euler angles *α* and *β* (rotation around x and y axis) expressing the tilt variations of the unaligned particles with respect to each other. Both histograms fit a normal distribution with a standard deviation of 14°.

**Supplementary Figure 5.**
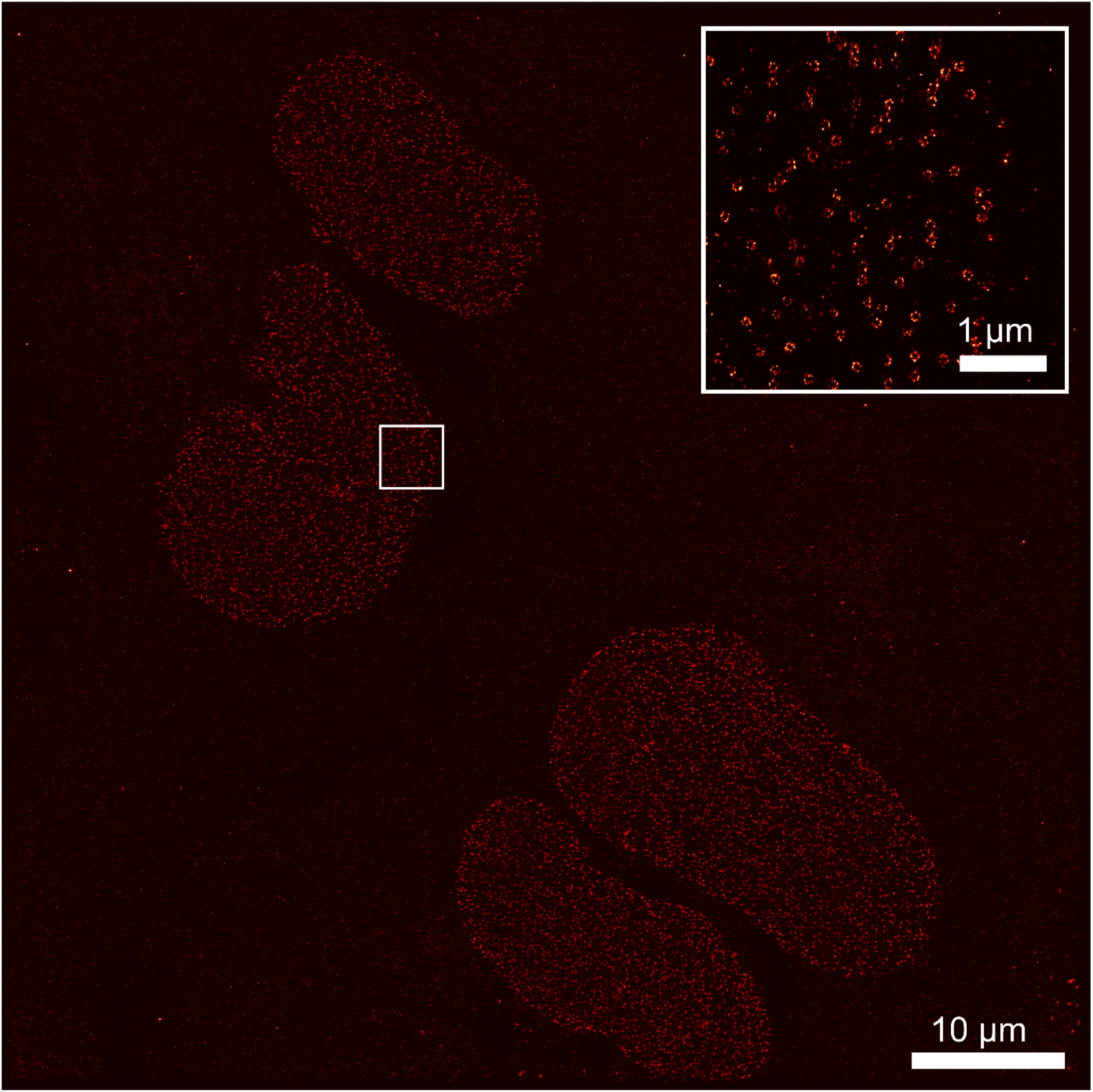
Whole field of view of SNAP-Tag labelled NUP107 proteins for DNA-PAINT imaging. The field of view shows the nuclei of four U2OS cells. The insert in the top-right corner presents a zoom in into the highlighted area.

**Supplementary Figure 6.**
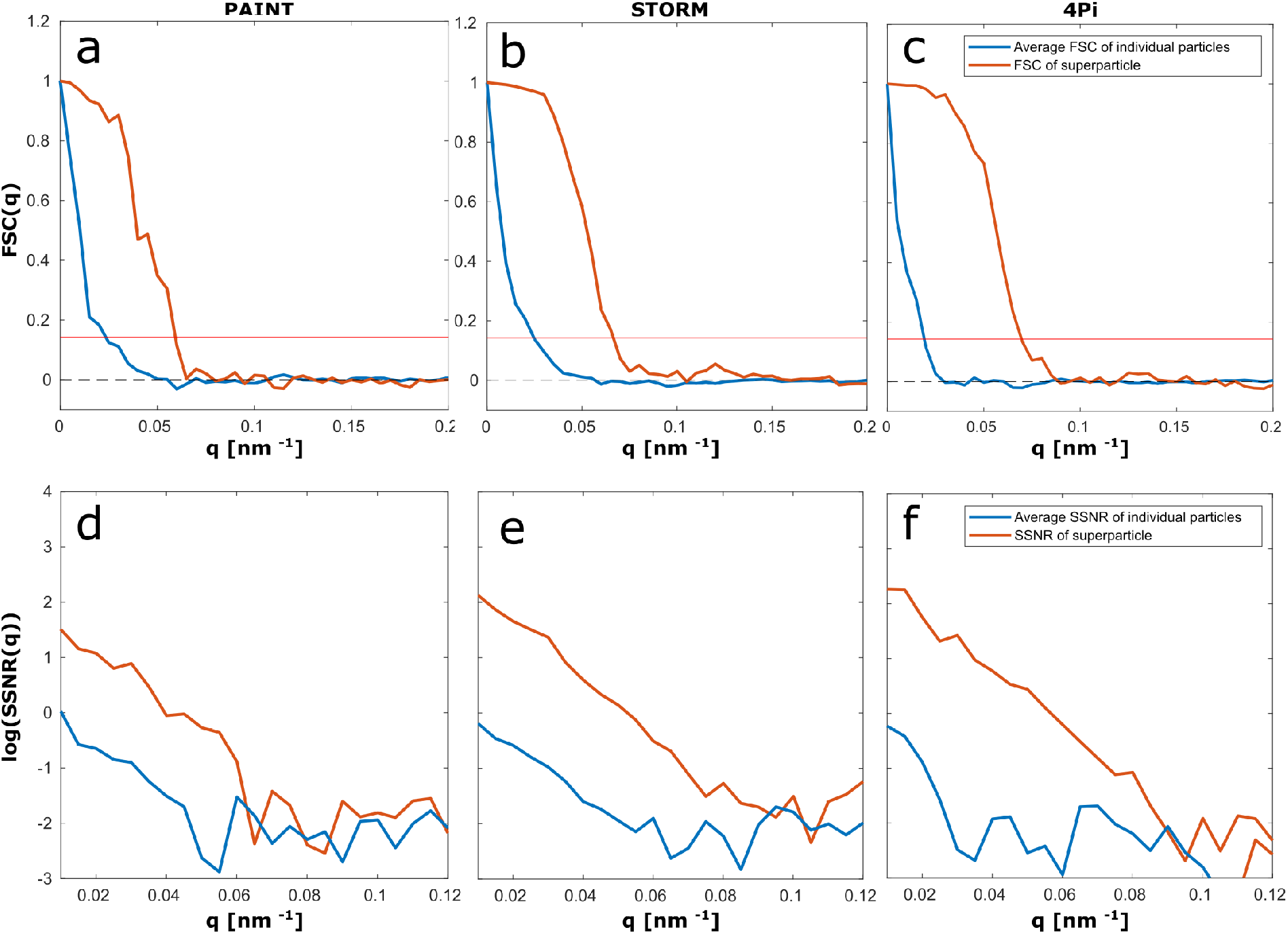
Fourier shell correlation^1^ (FSC) and spectral signal-to-noise ratio (SSNR) curves for the initial particles and the corresponding super-particles of 3D astigmatic PAINT, 3D astigmatic STORM and 4Pi STORM data. (**a-c**) The FSC curves show the resolution improvement from 42.6, 40.5, and 52.2 nm to 16.6, 15.1 and 14.2 nm for the three reconstructions respectively. (**d-f**) The SSNR curves show about two orders of magnitude improvement in spectral signal-to-noise ratio over all. These values are in good accordance with the visual quality of the super-particles. From these FSC values it is also clear that the dimers cannot be resolved which are at 12 nm distance according to the EM model. The FSC/SSNR curves for individual particles averages (blue) are computed between pairs of individual particles and then averaged. SSNR is computed as follows SSNR = FSC/(1−FSC).

**Supplementary Figure 7.**
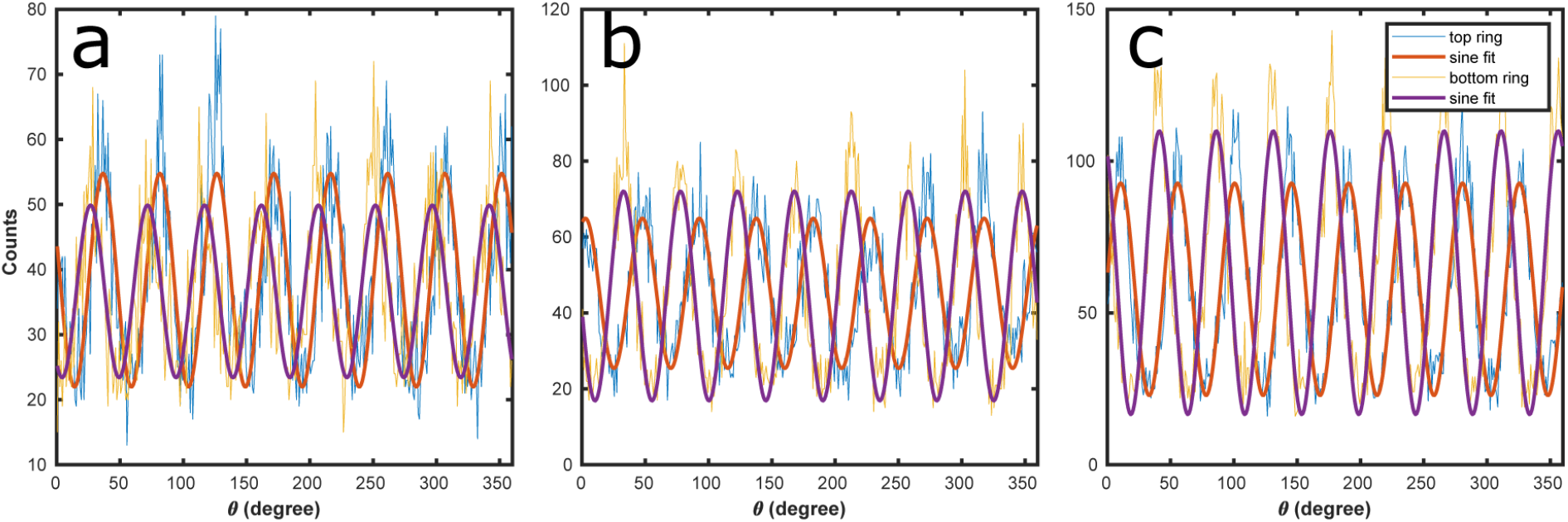
The distribution of the localizations over azimuthal angle and the fitted sine function for the super-particles in **Figure 2**. **(a)** PAINT reconstruction. **(b)** STORM reconstruction. **(c)** 4Pi reconstruction. In order to find the phase shift between the cytoplasmic and nuclear rings, we fit a sine function to the azimuthal angles of the localization data points in each ring. The difference in the phases of the fitted sine function for each reconstruction defines the azimuthal phase shift of the two rings.

**Supplementary Figure 8.**
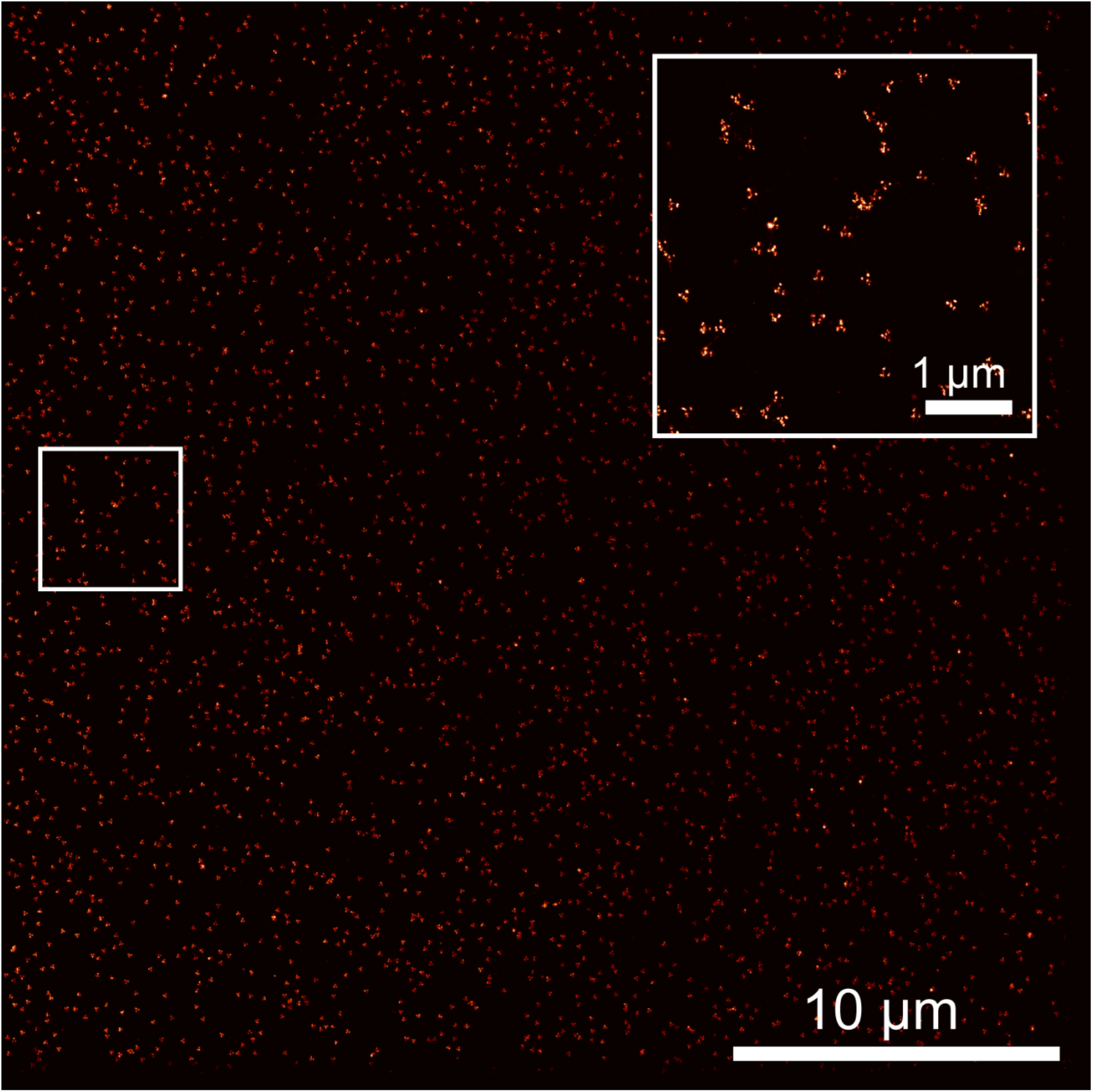
Whole field of view of three-dimensional DNA origami tetrahedron structures imaged with DNA-PAINT on a spinning disk microscope. The side length of the symmetric tetrahedron structure is 100 nm. The insert in the top-right corner presents a zoom in into the highlighted area.

**Supplementary Figure 9.**
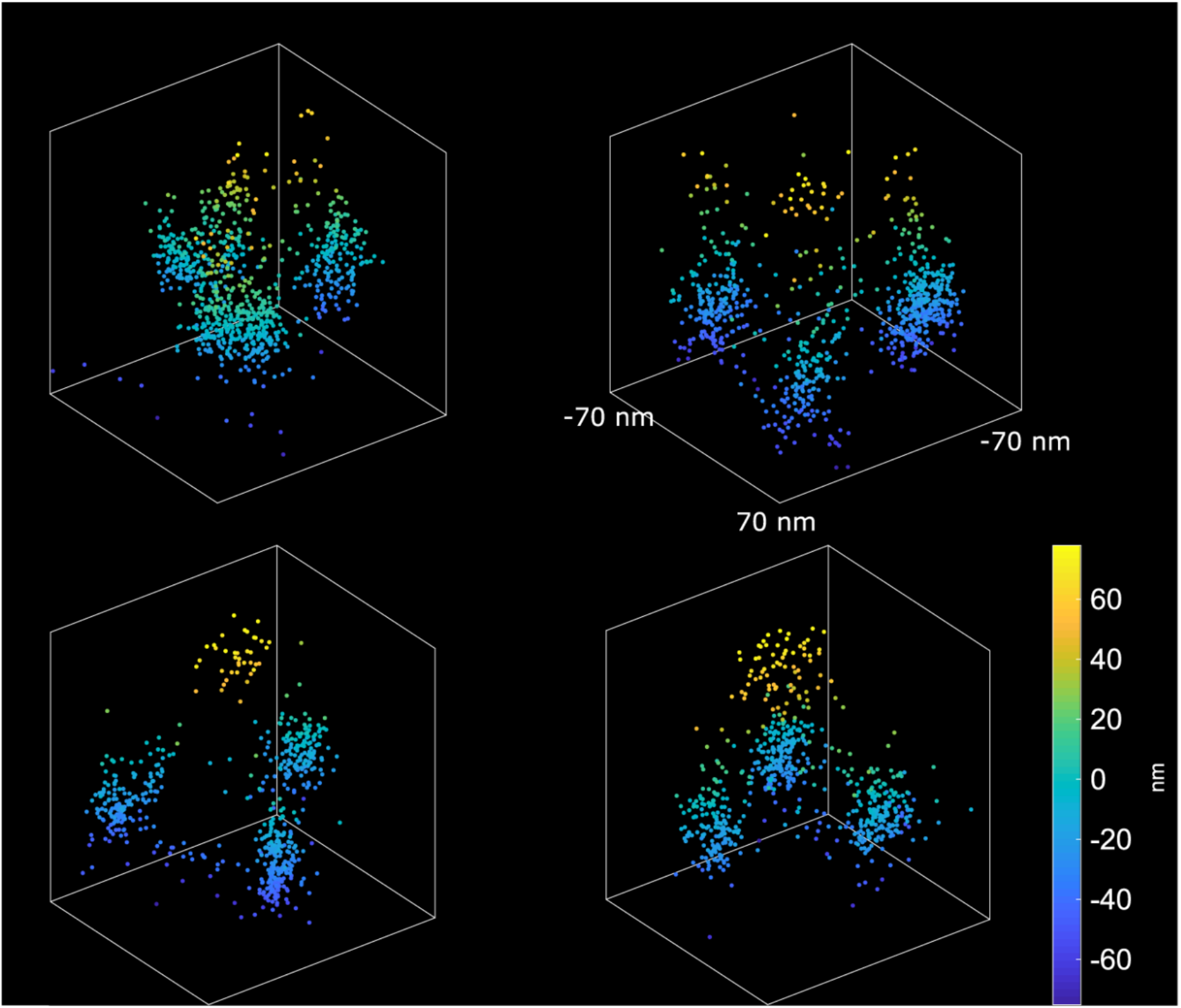
Example images of tetrahedron DNA-origami nanostructures imaged with PAINT.

**Supplementary Figure 10.**
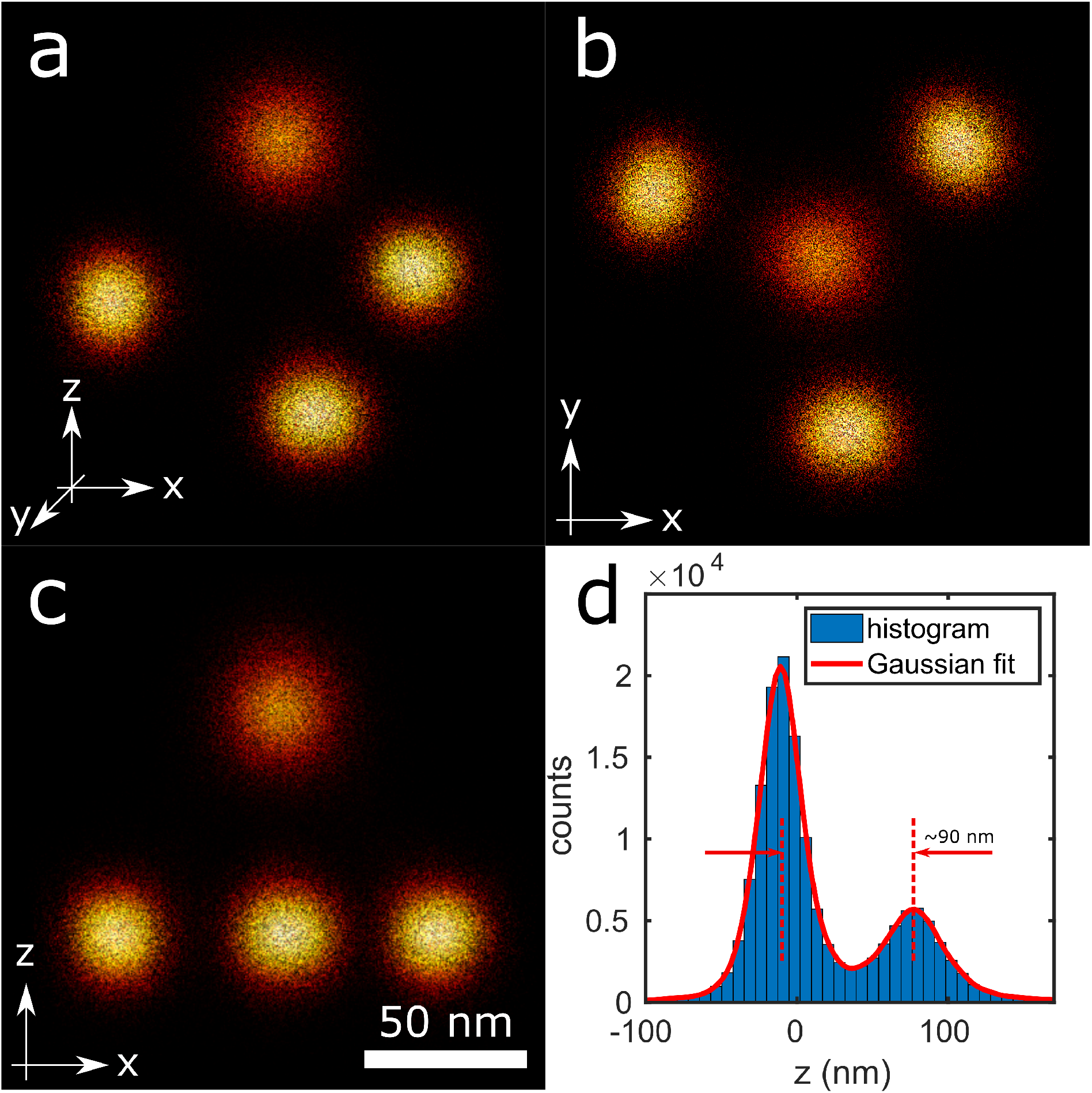
Fusion of 256 tetrahedron DNA-origami nanostructures. **(a)** Side view of the super-particle. **(b)** Top (x-y) view of the super-particle. **(c)** Front (x-z) view of the super-particle. **(d)** Histogram of the z coordinate of the localization data showing a distance of ~90 nm between the two peaks. Particle fusion of the nanostructures result in an isotropic distribution of the localization over the four binding sites of the tetrahedron as seen from the round localization distributions around the binding sites.

**Supplementary Figure 11.**
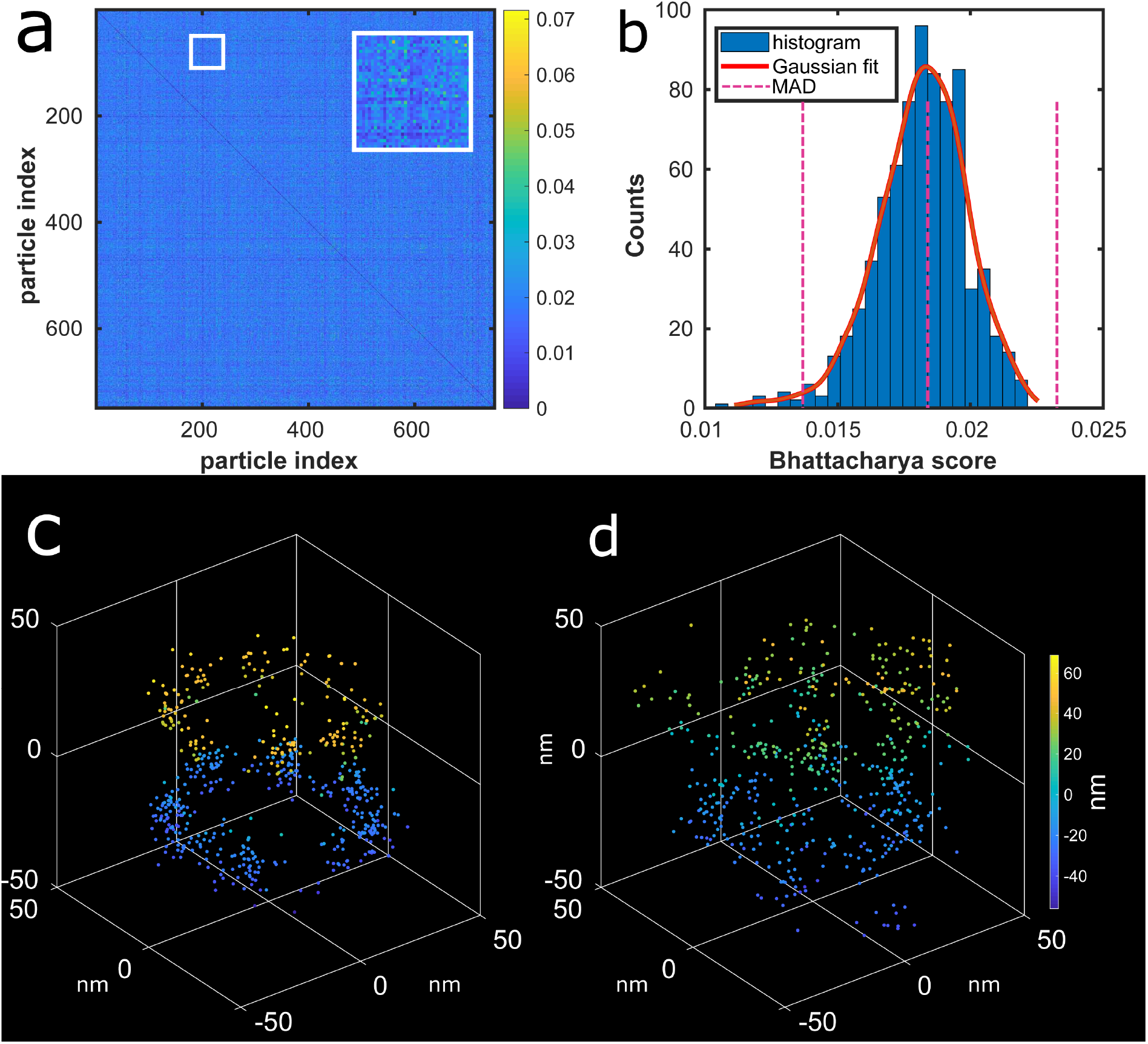
Outlier particle removal. **(a)** All-to-all Bhattacharya heat map matrix for the fusion of 750 particles from the 4Pi dataset. Each value is rendered as a pixel in a 750 × 750 image. **(b)** The histogram of the Bhattacharya scores (cost function value) of each particle which is obtained by averaging the matrix in **a** along the columns (or rows) together with the median absolute deviations (MAD) magenta line magenta and its lower and upper bounds. Only 9 particles are recognized as outliers in this dataset with these default settings of the MAD threshold. **(c)** Overlay of the localizations of the best particles (20 top scores). **(d)** Overlay of the localizations of the worst particles (20 lowest scores). While the (good) particles in **c** form a sharp super-particle, the overlay in **d** is quite blurry and the localizations are more scattered around the NUP structure.

**Supplementary Figure 12.**
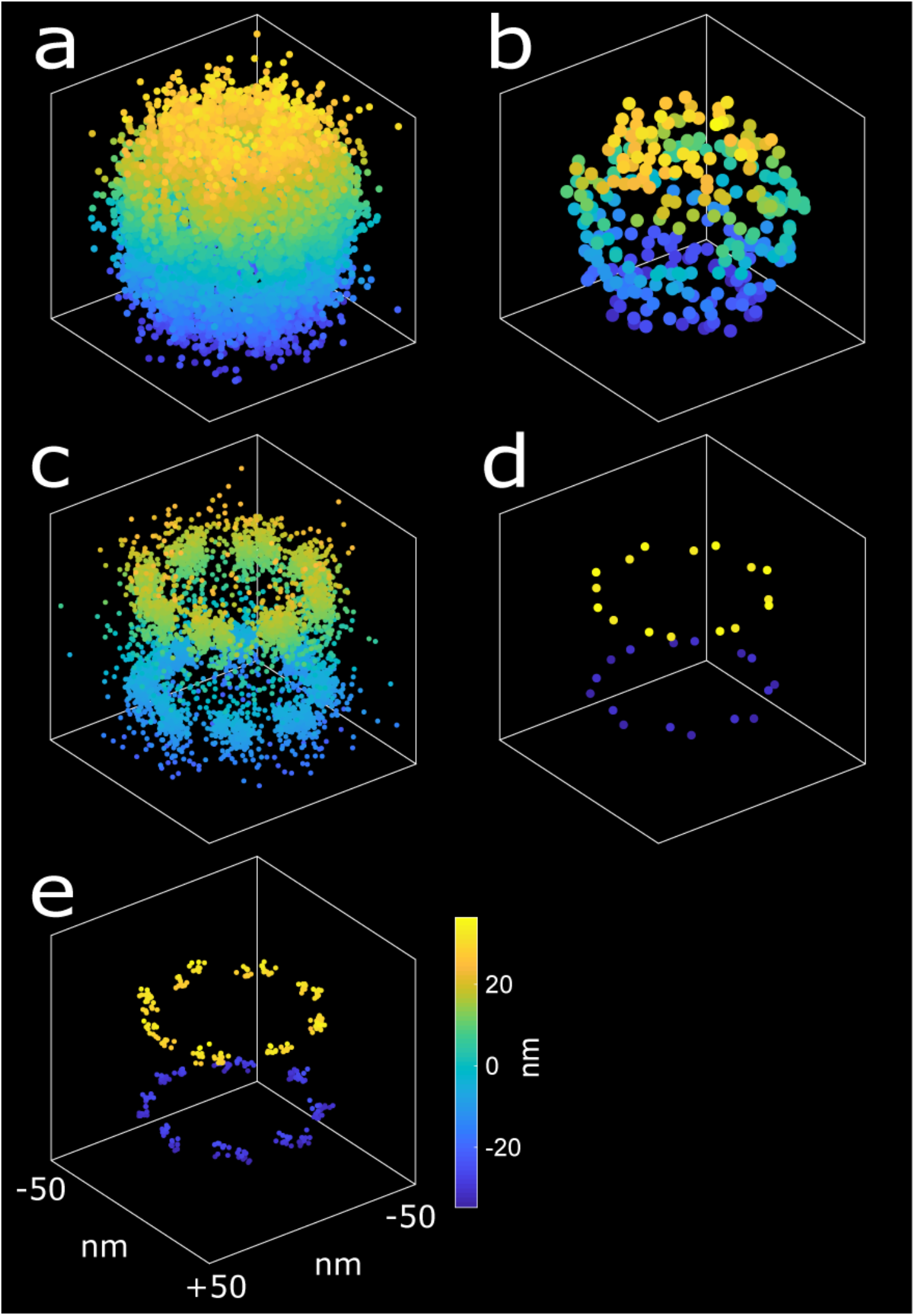
The principle of the proposed registration error measurement. **(a)** Overlay of 10 simulated particles before alignment. **(b)** Overlay of the binding sites of the particles in **a**. **(c)** Superparticle as a result of fusing the particles in **a**. **(d)** The corresponding binding sites of the aligned particles in **c** in a perfect fusion (zero measurement error). **(e)** The corresponding binding sites of the aligned particles in **c** with the effect of the registration error taken into account. Ideally and in a perfect fusion, all the binding sites of the ground-truth simulation model should co-locate. Due to the registration error they scatter around the mean shape model of the super-particle. The corresponding registration error for a run of the particle fusion pipeline is found by quantifying this scatter.

**Supplementary Figure 13.**
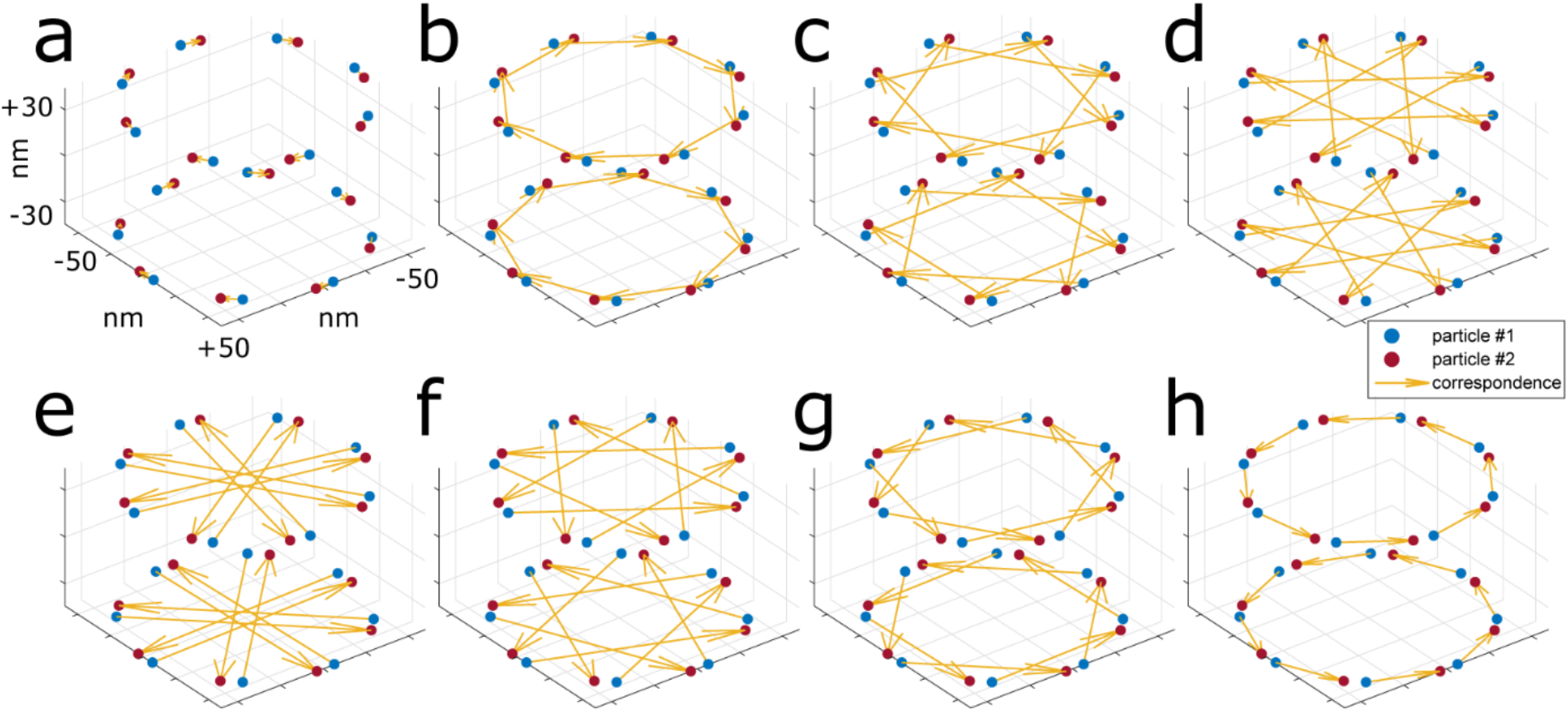
Correspondence problem in matching binding sites of two aligned particles. **(a-h)** Different correspondence possibilities for computing the error between the registered particles with registration error. In this example, each particle includes 16 binding sites. Since the binding sites are ordered, there are only 8 different combinations of the correspondences between them. The minimum Euclidean distance among these eight candidates defines the correct correspondence and its value is the alignment error.

**Supplementary Figure 14.**
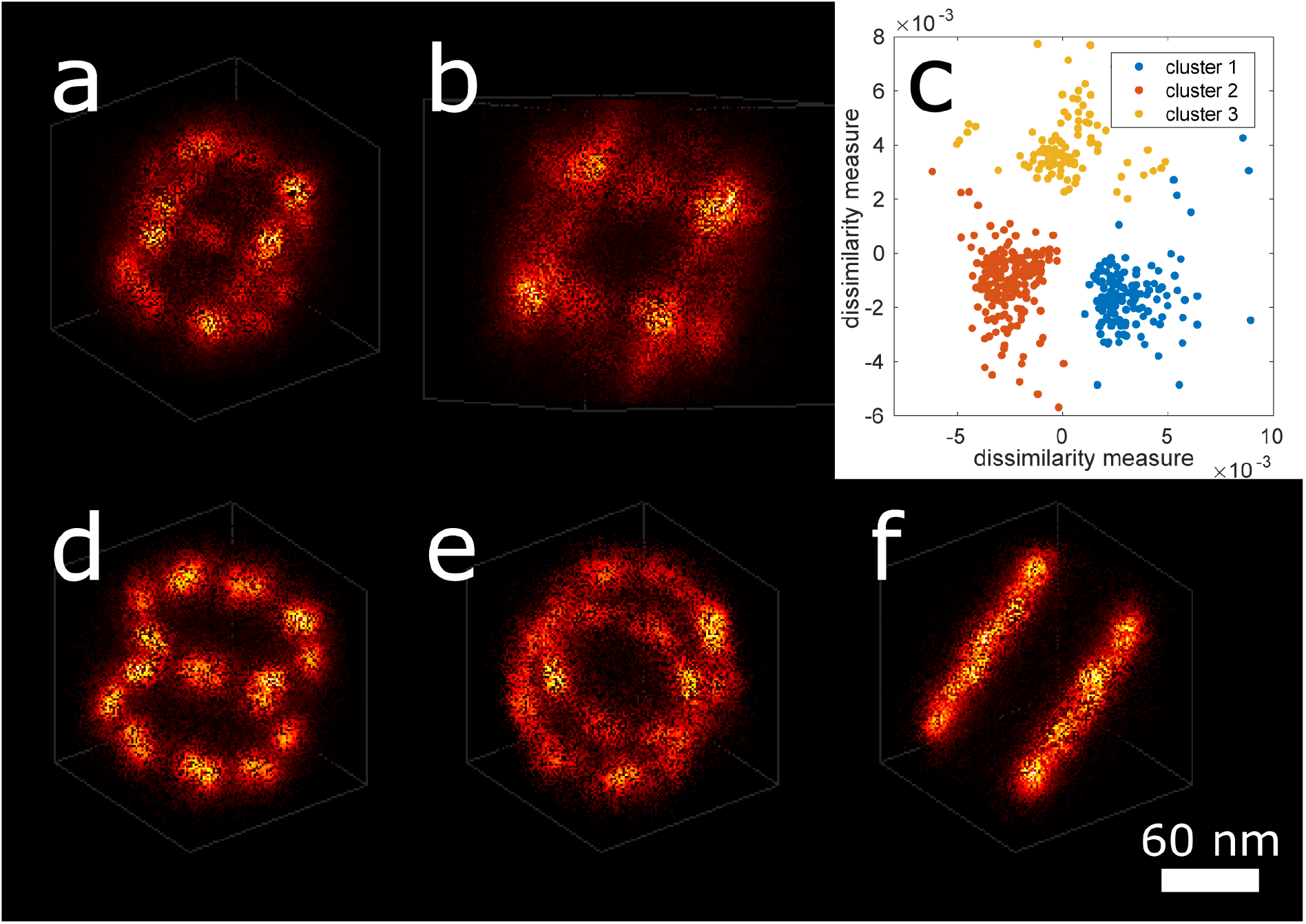
Fusion of particles with arbitrary poses. **(a-b)** Two views of the initial super-particle after the bootstrapping step for fusing 400 simulated PAINT particles. **(c)** K-means clustering (*k* = 3) on multi-dimensional scaling of the dissimilarity matrix of the all-to-all matrix. **(d-f)** Three clusters of particles which are separated using the proposed method each containing 176, 96 and 128 particles, respectively. When the initial particles have arbitrary poses, the particle fusion results in clusters of particles which are aligned together. To separate these clusters, the all-to-all Bhattacharya score matrix can be used to map them to the two-dimensional Cartesian space using multi-dimensional scaling (MDS). By clustering the MDS, particles which are aligned together can be automatically separated.

